## Supplementary figures for "Emergence of syntax and word prediction in an artificial neural circuit of the cerebellum"

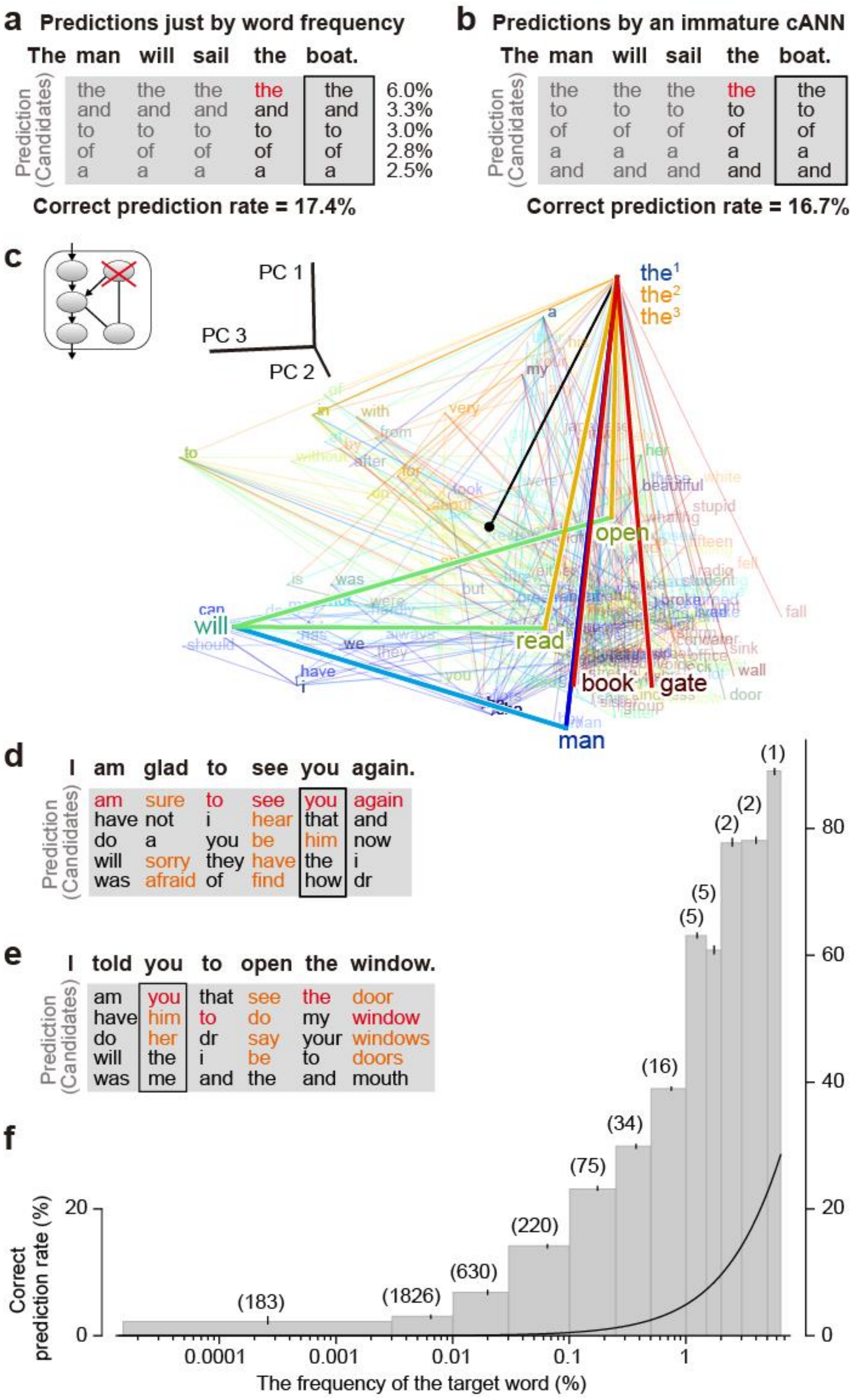


**Extended Data Figure 1.** Prediction and network dynamics of cANNs. **a**, Predictions just by the frequency of word occurrence (top 5 words). The frequency (%) for each word is indicated in the far-right column. **b**, Prediction of words (five candidate words), similar to that in **a**, by an immature cANN at a very early stage of learning (Fig. 1d, gray arrowhead). **c**, Neuronal activity in Purkinje cells after input of the$^1$, the$^2$, and the$^3$ in the two sentences from Figure 2a,b when the recurrent signal was blocked in the well-trained cANN of Figure 2. The dynamics of neuronal activity were visualized by PCA. **d,e**, Predictions of objective case pronouns after verbs (black rectangles). Correct predictions are highlighted in red. **f**, Correct prediction rate (%) for each group by frequency of occurrence of the target word (%). The number of words for each group is indicated in parentheses. The chance level (black trace) was calculated as $1-(1-p)^5$ to compare with the correct prediction rate of the five candidates.

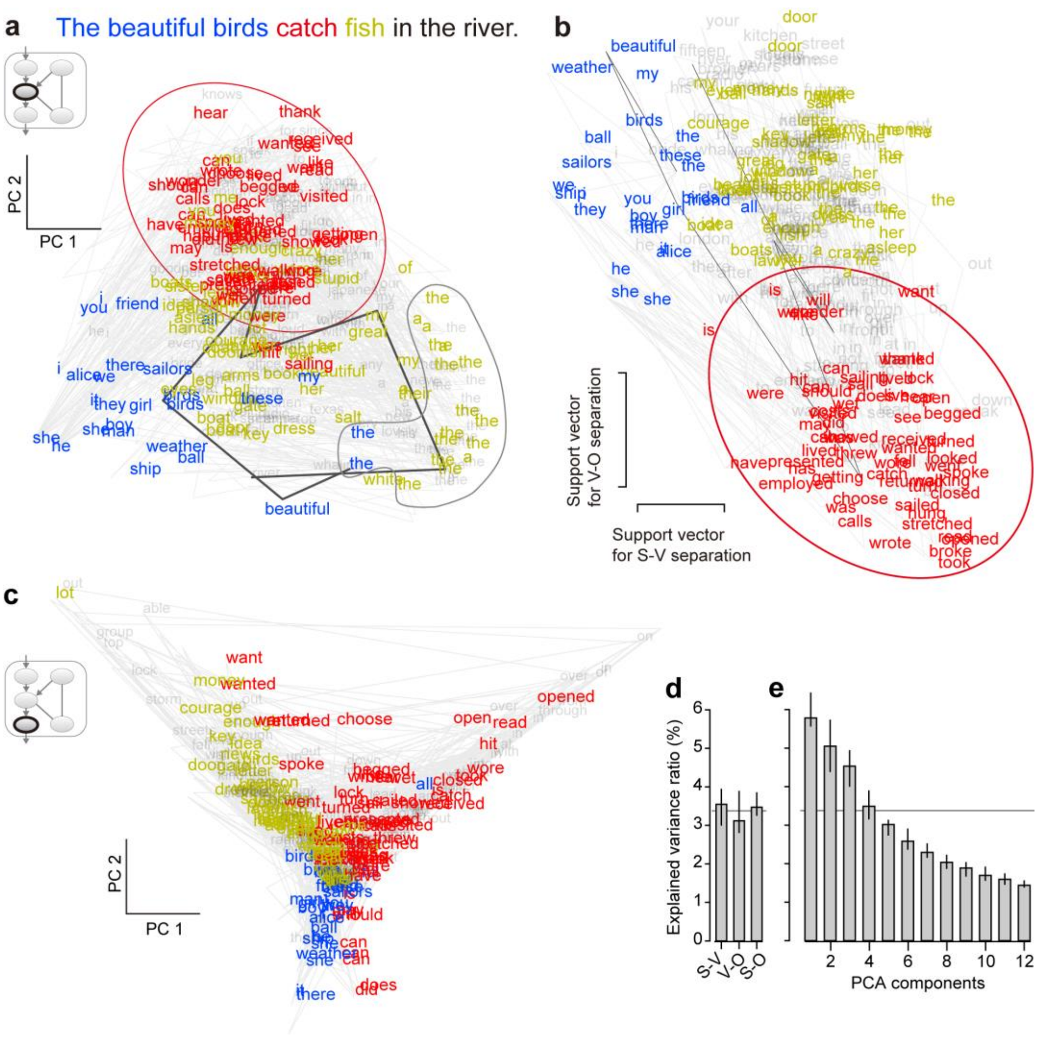


**Extended Data Figure 2.** Syntactic processing by the cANN. **a**, Neuronal activity of Purkinje cells in the cANN of Figure 2 after input of a subject (blue), verb (red), and object (yellow), visualized by PCA. The dynamics of activity for "The beautiful birds catch fish in the river" is indicated by the solid black trace. The verb cluster (red) is highlighted with an ellipse. The Purkinje cell signal clustered for the article words ('a' and 'the') separately from other words (gray border), suggesting the potential ability of word class recognition. **b**, Visualization of neuronal activity by the dimensions of the S-V and V-O classifications of the support vector machine. **c**, Neuronal activity of the output cells after input of a subject (blue), verb (red), and object (yellow), visualized by PCA. **d**, Percentage (median and interquartile range) of information (variance) that can be represented by the classification dimensions of the support vector machine over the total information (192 dimensions) for the 20 cANNs. **e**, Percentage of information (variance) that each PCA dimension can represent over the total information (192 dimensions). For comparison, the average of the three support vector dimensions in **d** is shown by the solid gray line. Note that the dimensions generated by the linear support vector machine represent about 3% of the total information, comparable to major dimensions of PCA space (e.g. the fourth).

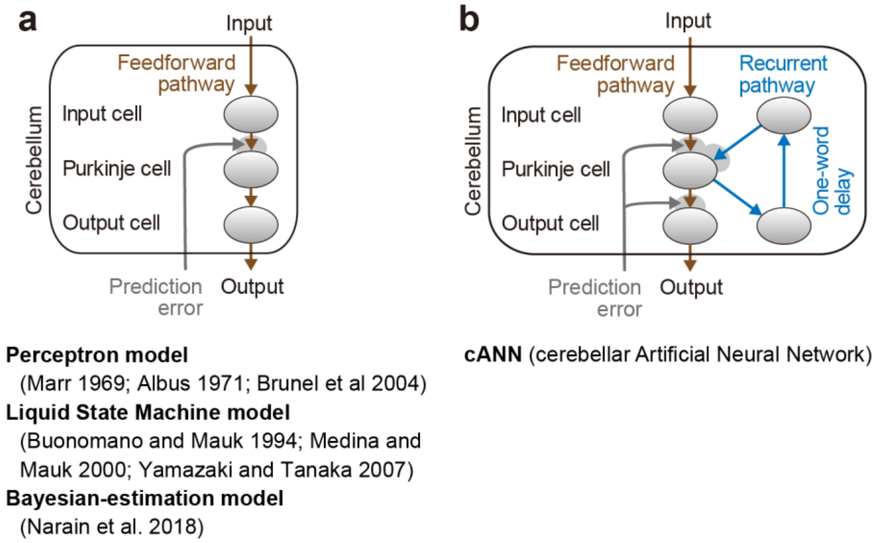


**Extended Data Figure 3.** Comparison of the cANN with conventional circuit models of the cerebellum. **a**, Conventional cerebellar circuit models, which contain only the feedforward pathway, have been proposed primarily to explain motor functions and, to our knowledge, have not yet been extended to language functions. The conventional models were prototyped from the perceptron model (Marr 1969; Albus 1971; Brunel et al 2004), which evolved into the Liquid State Machine model, which is a powerful sequence recording and reproducing machine (Buonomano and Mauk 1994; Medina and Mauk 2000; Yamazaki and Tanaka 2007). These are suitable for explaining motor functions in affinity with the internal-model theory of the cerebellum. In addition, there is an advanced version of the model equipped with Bayesian estimation (Narain et al. 2018). It would be possible to extend these to predict the next word, but it would be difficult to output syntactic information. This is because all the conventional models based on the perceptron model are two-layer neural networks with Purkinje cells as the output layer and no intermediate layer (cerebellar nucleus neurons are largely responsible for relaying information from Purkinje cells). Therefore, there is no layer corresponding to the intermediate layer of the cANN which extracts the syntactic information. In addition, although the evolved Liquid State Machine models and later models have an input layer that can encode more dynamic and complex information than the input layer of the cANN, it is invariant before and after learning and cannot learn syntactic information. **b**, The cANN contains a recurrent pathway, and it is new because it can not only predict the next word in the output layer but also extract syntactic information in the intermediate layer, unifying the two language functions and better explaining the findings of language disorders of the cerebellum.

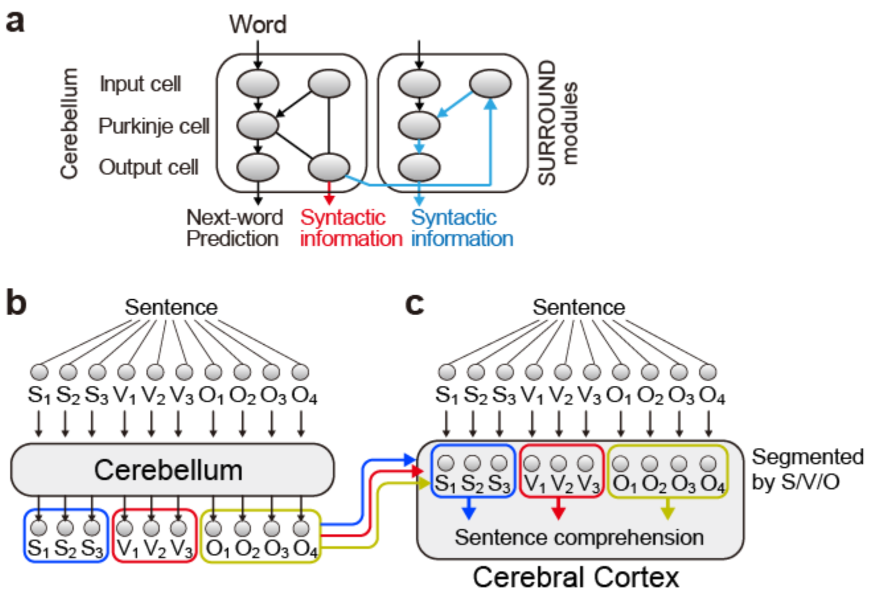


**Extended Data Figure 4.** Output channel for syntactic information and neocortico-cerebellar roles, proposed by the cANN model. **a**, Candidates for the output channel for the syntactic S-V-O information. Since the Purkinje cells send the syntactic information to the recurrent pathway, if information can be sent out from any part of this pathway, the cANN could function as a syntax-processing circuit. Considering that the cerebellum has a modular structure, there are two candidates for the output channel. The first possibility is from the recurrent-output neurons in the next-word prediction module (red arrow). Considering the spread of the recurrent signal from the output layer of the next-word prediction module to the input layer of the neighboring modules (Apps and Garwicz, 2005; Gao et al., 2016; Ohmae et al., 2021), the second possibility is through the input layer–Purkinje cell–output layer of a neighboring module (blue pathway). This pathway can form a parallel but different output channel from the main prediction output (Chen & Evinger 2006; Ohmae et al. 2021). **b**, Non-hierarchical sentence processing in the cerebellum. The cerebellum receives unprocessed raw words and directly processes the sequence. Because it often needs to process long sequences, processing may require an enormous number of granule cells (i.e., input cells in the cANN) to retain word history (cerebellar granule cells occupy more than half of the total number of neurons in the entire brain). **c**, Hierarchical sentence processing in the neocortex. Since the neocortex cannot process sequences as long as the cerebellum, the neocortex preprocesses the sentence by grouping the words into subjects, verbs, and objects to generate sentence segmentations, so that the neocortex only needs to process shorter sequence. We propose that, during development, this segmentation process in the neocortex requires syntactic information from the cerebellum.
